# Protective effect of peroxisome proliferator-activated receptor-α in regulating the early inflammation response to extended hepatectomy from a comparative transcriptomic analysis

**DOI:** 10.1101/2021.12.12.472295

**Authors:** Cheng-Cheng Shi, Yang Bai, Xin Yan, Nuo Cheng, Wen-Zhi Guo, Shui-Jun Zhang, Ji-Hua Shi

## Abstract

The regulation mechanism of small-for-size syndrome remains unclear. Thus, we aimed to analyze the molecular profiles following extended hepatectomy and identify the therapeutic target. Major hepatectomy and extended hepatectomy were performed in the rat model, and the remnant livers were obtained dynamically for the high-throughput transcriptome analysis to identify the differentially expressed genes (DEGs). The general framework for weighted gene co-expression network analysis (WGCNA) was employed to explore the expression patterns of DEGs. As result, WGCNA identified 10 distinct gene co-expression modules according to the correlation between module eigengene and different postoperative time-points. The magenta module (gene count: 289) and the lightcyan module (gene count: 484) were found positively correlated with major hepatectomy instead of extended hepatectomy. In the lightcyan module, peroxisome proliferator-activated receptor-α (PPARα) was selected and found the down-regulation in the remnant liver following extended (marginal) hepatectomy in rats and humans. Besides, administration of PPARα agonist attenuated hepatic inflammation injury while PPARα antagonist increased liver inflammation injury after extended hepatectomy in rats, marked by the significantly changed aminotransferases, tumor necrosis factor-α and interleukin*-*6 levels in the plasm, and histological Suzuki criteria. Consequently, DEGs and their molecular profiles after extended hepatectomy were identified, and PPARα might be a potential therapy target for small-for-size syndrome.

## Introduction

Small-for-size syndrome (SFSS), which is essentially post-operative hepatic failure, may occur after extended hepatectomy and reduced-size liver transplantation. Pathogenesis and clinical manifestations of SFSS are characterized with portal hypertension, inadequate liver function and compromised liver regeneration accounting for a high mortality rate (Riddiough et al. 2020; Rajakumar et al. 2017). In order to accelerate the postoperative parenchymal restoration and thus reduce the mortality rate, most of previous studies have focused on the mechanism underlying portal hyperperfusion and the suitable size of remnant liver (Gilgenkrantz and Collin de l’Hortet 2018; Rajakumar et al. 2017; Wong et al. 2021). Portal hypertension may cause severe liver damage and reduce parenchymal restoration through over-release of inflammatory cytokines (Athanasiou et al. 2017; Govil 2020). A recent study indicated that the acute kidney injury is associated with extended hepatectomy, and a sufficient future liver remnant could avoid postoperative injury (Reese et al. 2021). The efforts to explore the mechanism of the systemic inflammation and attenuate the liver injury of the remnant liver are still limited (Riddiough et al. 2020; Rajakumar et al. 2017).

Sequencing technology and mass spectrometry-based proteomics create the superior research platforms for the further molecular mechanism. Wu JD et al (Wu et al. 2010) characterized the protein expression patterns during the early phases of a rat model using proteomics, and identified potential regulators by functional analysis of these SFSS-related proteins, which still require further verification. The regulation networks following surgery are too complex to outline clearly so far. Thus, the underlying mechanisms of SFSS regulation still require more efforts.

Sequencing-based transcriptome detection allows enhanced detection of low-abundance transcripts and proteomics, and the further bioinformatics analysis could provide analytical information on expression and regulation of genes, and may be predictive for revealing the mechanism of the molecular response (Lopes et al. 2011). The general framework for weighted gene co-expression network analysis (WGCNA) was employed to find co-expressed gene modules with hub genes and define the gene expression network topology (Zhang and Horvath 2005). The constructed co-expression modules usually correspond to the specific biological pathways (MacLennan et al. 2009), and the further analysis could shed light on the underlying regulation mechanism. The purposes of the current study were to explore the regulation profiles of SFSS by using next-sequencing sequencing and WGCNA, and thus to select the potential therapeutic targets for the clinical usage.

## Materials and Methods

### Clinical study of the patients after partial hepatectomy

The human study protocol was approved by the Research Committee of the Affiliated Hospital of Zhengzhou University (registration number: 2019-KY-21) and conformed to the ethical guidelines of the 1975 Declaration of Helsinki (Puri et al. 2009). A number of 12 patients were included in the study, which were undergone open partial hepatectomy with liver cancer and obtained with the informed consent. The clinico-pathological characteristics are summarized in Supplemental Table S1. According to its resection size, the patients were grouped into marginal hepatectomy with resection size over 45% (Group M, N=8) and minor hepatectomy (wedge and segment resection) with resection size less than 30% (Group C, N=6). The distant non-tumor liver tissue was obtained at one hour (1 h) post-surgery to verify the selected targets from the preclinical analysis in rats. We used no organ from the executed prisoners but the samples from the patients after partial hepatectomy.

### Development of partial hepatectomy in rats

The animal experimental protocol was supervised and approved by the ethics committee for animal experiments of Zhengzhou University (2019-KY-183), and was performed in accordance with the ARRIVE guidelines. The male Sprague-Dawley rats with 12-week-old (Keli Experimental Animal Center, Beijing, China) from Henan Experimental Animal Center in Zhengzhou University were used.

The extended hepatectomy with the resection volume between 70% and 90% in rats is most widely used to model SFSS following either extended hepatectomy or reduced-size liver transplantation (Nacif et al. 2015; Dahmen et al. 2008; Wakabayashi et al. 2021). Based on our previous study, 80% hepatectomy compromises liver function in rats without conferring mortality risk compared with 90% hepatectomy (Shi et al. 2011). Thus, we employed 80% hepatectomy to simulate SFSS. Hepatectomy with resection size of 70% was used as the standard regeneration model to compare the regenerative capability with 80% hepatectomy (extended hepatectomy). A modification of the partial hepatectomy (PH) technique was used to reduce the liver size *in situ* under anesthesia (Shi et al. 2011).

The animal experiment was grouped into two cohorts. In the detection cohort, as reported in the earlier publication (Yan et al. 2019), according to the degree of regeneration stimulation by its resection size, a total of 33 rats were grouped into Group PH70 (70% major hepatectomy, N=15), Group PH80 (80% extended hepatectomy, N=15) and control (sham control, N=3) (Table 1). A number of 3 rats in each group were euthanized to harvest blood and tissue samples at 1h, 6h, 12h, 24h, 72h after partial hepatectomy. The sham-operated rats were regarded as control, whose tissue were harvested. Liver total RNA was isolated with TRIzol reagent (Invitrogen, Carlsbad, CA) for the further high-throughput sequencing detection, and the RNA concentration was measured using Nanodrop2000 (Thermo Fischer, Waltham, USA).

**Table 1.** animal group and treatment.

|  | Group | Hepatectomy | Treatment | Time point |  |  |  |  |  |
| --- | --- | --- | --- | --- | --- | --- | --- | --- | --- |
|  |  |  |  | before<br>hepatectomy | 1h | 6h | 12h | 24h | 72h |
| **The**<br>**detection**<br>**cohort** | **Control** | - | - | n=3 |  |  |  |  |  |
|  | **PH70** | 70% hepatectomy | - |  | n=3+3 | n=3+3 | n=3 | n=3 | n=3 |
|  | **PH80** | 80% hepatectomy | - |  | n=3+3 | n=3+3 | n=3 | n=3 | n=3 |
| **The**<br>**verification**<br>**cohort** | **DMSO** | 80% hepatectomy | 10% DMSO,<br>IV* |  | n=3 | n=3 |  | n=3 | n=3 |
|  | **WY14643** | 80% hepatectomy | WY14643 in<br>10% DMSO,<br>IV* |  | n=3 | n=3 |  | n=3 | n=3 |
|  | **GW6471** | 80% hepatectomy | GW6471 in<br>10% DMSO,<br>IV* |  | n=3 | n=3 |  | n=3 | n=3 |

\*DMSO, dimethyl sulfoxide; IV, intravenous injection.

The liver tissues, obtained at specific time points following 70% major hepatectomy (N=6) and 80% extended hepatectomy (N=6) in rats, were used to ascertain the expressions of the differentially expressed genes by quantitative reverse transcription polymerase chain reaction (RT-qPCR) and western blot were used to detect the gene expression of the early response to partial hepatectomy (Table 1).

In the verification cohort, in order to verify the function of the differentially expressed gene from the detection cohort, 36 rats with 80% hepatectomy were grouped into Group DMSO (80% extended hepatectomy +10% dimethyl sulfoxide intravenous injection, N=12), Group WY14643 (80% extended hepatectomy + postoperative intravenous injection of PPARα agonist WY14643, N=12), and Group GW6471 (80% extended hepatectomy + postoperative intravenous injection of PPARα antagonist GW6471, N=12) (Table 1). A number of 3 rats in each group were euthanized to obtain plasm and tissue samples at 1h, 6h, 24h and 72h after partial hepatectomy. Liver regeneration was evaluated by the ratio of remnant liver weight relative to body weight (LBW) (Shi et al. 2017). The sham-operated rats, whose tissue were obtained before surgery (0h), were regarded as control. Plasm samples were obtained from the abdominal aorta to measure the levels of alanine aminotransferase (ALT), aspartate aminotransferase (AST), tumor necrosis factor-α (TNF-α) and interleukin-6 (IL-6). Liver samples from the remnant livers were obtained to detect the expression of hub gene and the assess parameters of hepatic injury.

### Transcriptome sequencing

Total RNA from rat liver samples was extracted according to the manual of mirVana mRNA Isolation reagent (Ambion, USA). The transcriptome sequencing was performed in collaboration with OE Biotech Co., Ltd. (Shanghai, China) using the Illumina Hiseq X Ten platform (Illumina Inc., San Diego, USA). Fragments per kilobase million and read counts value of each transcript were calculated by Cufflinks (Trapnell et al. 2010) and HTSeq-count (Anders et al. 2015) respectively. The project and the raw data of the transcriptome sequencing have been registered and uploaded in the National Center for Biotechnical Information (PRJNA687124). The fold change and *P* value were calculated by DESeq R package to obtain the differentially expressed genes. Differentially expressed genes were determined by the criteria of fold change >2 with a *P* value < 0.05. Venn diagram was drawn using the VENNY software (http://bioinfogp.cnb.csic.es/tools/venny).

### WGCNA

The co-expression network analysis was performed with the Pearson correlations between all gene pairs, then converted the correlation matrix into an adjacency matrix using the WGCNA package in R (Version 1.69) (Zhang and Horvath 2005). WGCNA employed a soft threshold based on the determination method (MacLennan et al. 2009). The parameter of the power function was determined by which the resulting adjacency matrix was transformed into topological overlap matrix and the gene-network connectivity was defined as the sum of its adjacency with all other genes for network generation. A positive correlation (*P*<0.05 and r>0.30) was set to select to unravel the biological pathways underlying the traits. Genes with high intramodular connectivity are considered as intramodular hub genes.

### Gene ontology (GO) and Kyoto Encyclopedia of Genes and Genomes (KEGG) pathway enrichment analysis

After construction of co-expression modules from WGCNA, GO and KEGG pathway analysis of the differential expressed genes related to biological functions were performed respectively using R package based on the hypergeometric distribution as prescribed before (Kanehisa et al. 2008). The *P* <0.05 was considered significantly enriched for GO and KEGG pathway.

### RT-qPCR

RT-qPCR was performed as previously described (Shi et al. 2014). The gene related to key modules of extended hepatectomy, peroxisome proliferator-activated receptor-α (PPARα), was selected for RT-qPCR assays to verify the transcriptome sequencing and WGCNA. The primers of PPARα and PPARα-downstream regulated gene cytochrome P450 1A2 (CYP1A2) (Pant et al. 2019) were designed and synthesized by Shanghai Generay Biotech Co., Ltd (Supplemental file 2). Quantitative analysis of gene expression was performed on ABI Prism 7000 (Applied Biosystems, Foster City, CA) using 2× SYBR Green master mix (Applied Biosystems, Foster City, CA). Gene expression was presented relative to the level of housekeeping gene glyceraldehyde-3-phosphate dehydrogenase (GAPDH, Supplemental Table S2).

### Western blot

Protein extracts of liver tissue were prepared and processed for western blot analysis as previously described (Shi et al. 2011; Shi et al. 2020). Primary antibodies used were anti-PPARα (1:1000, 15540-1-AP, Proteintech, Rosemont, USA), CYP1A2 (1:1000, 19936-1-AP, Proteintech, Rosemont, USA) and GAPDH (1:5000, 60004-1-Ig, Proteintech, Rosemont, USA). Immunoreactivities were visualized by secondary horseradish peroxidase-conjugated rabbit (1:2000, SA00001-2, Proteintech, Rosemont, USA) and mouse (1:2000, SA00001-1, Proteintech, Rosemont, USA).

### Hematoxylin and eosin (HE) staining and histological evaluation

Liver tissue were fixed in 4% paraformaldehyde in phosphate-buffered, embedded in paraffin wax, and stored at 4°C. HE staining was performed as before (Shi et al. 2011; Shi et al. 2020). Histological analysis was evaluated according to Suzuki histological criteria (Shi et al. 2020; Suzuki et al. 1993).

### Enzyme-linked immunosorbent assay (ELISA)

Inflammatory cytokine (TNF-α and IL-6) levels in the plasma were measured by ELISA according to the manufacturer’s protocols (KE20001, Proteintech, Wuhan, China; EK0412, Boster Biological Technology, Wuhan, China).

### Statistical Analysis

Values are expressed as means with standard deviation (SD). Differences in groups were analyzed by the Student’s test and one-way ANOVA. The statistical analysis was employed by SPSS 21.0 (IBM, Armonk, New York, USA). A probability level of less than 5 per cent (*P*<0.05) was considered statistically significant.

## Results

### Identification of the differentially expressed genes following hepatectomy

The genome-wide mRNA expression profiles in the liver biopsies after major hepatectomy and extended hepatectomy were detected. Compared with the control (the time-point of before surgery), 14170 and 14113 dysregulated genes for at least one time-point after major hepatectomy and extended hepatectomy were detected respectively (Fold Change >2 and *P* value < 0.05) and expressed in Figure 1. As shown in the Venn diagram (Figure 1A and 1B), it was found that 10319 and 10518 genes were significantly expressed at 1h, 10745 and 11218 genes at 6 h, 12072 and 11339 genes at 12 h, 10536 and 11135 genes at 24 h, 10613 and 10796 genes at 72 h after major hepatectomy and extended hepatectomy, respectively (Figure 1C). The expression pattern of the differentially expressed genes of major hepatectomy and extended hepatectomy was shown in Figure 1C.

**Figure 1.**
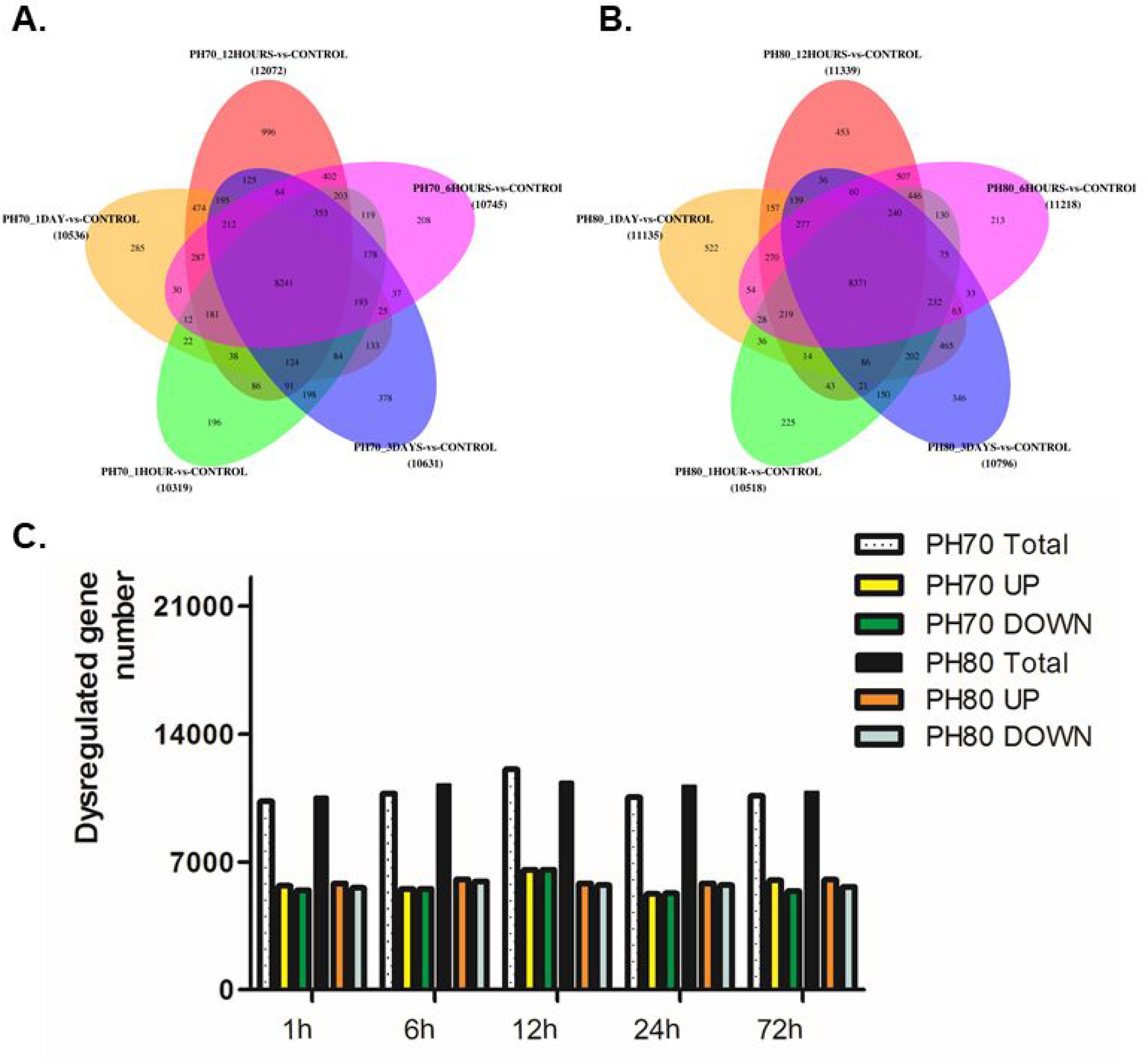
Expression of the differentially expressed genes related to the early response after major hepatectomy (PH70) and extended hepatectomy (PH80) in rats by transcriptome sequencing. The Venn diagram illustrates the overlap of differently expressed genes after PH70 (Figure 1A) and PH80 (Figure 1A) (Fold change >2.0, *P*<0.05); Figure 1C, columns represent the numbers of differently expressed genes after hepatectomy.

### Construction of WGCNA and identification of modules associated with extended hepatectomy

From 33 samples of major hepatectomy and extended hepatectomy in rats, expression values of 9220 differentially expressed genes were obtained and subjected to construct the co-expression modules of WGCNA.

We chose a soft-threshold power of 18 with a minimum module size 30 (check scale topology, power:18, scale R^2^=0.53, SLOPE=-0.89) to define the adjacency matrix. WGCNA identified a number of 10 distinct gene co-expression modules after hepatectomy (Figure 2A), and then intramodular connectivity (Figure 2B) and gene significance was used to identify hub genes for further validation. Interaction analysis identified the co-expression module associated with particular traits according to the correlation between module eigengene and time-points after hepatectomy (Figure 2C). The color code in the module-trait correlation is based on Pearson’s correlation value and *P* values.

**Figure 2.**
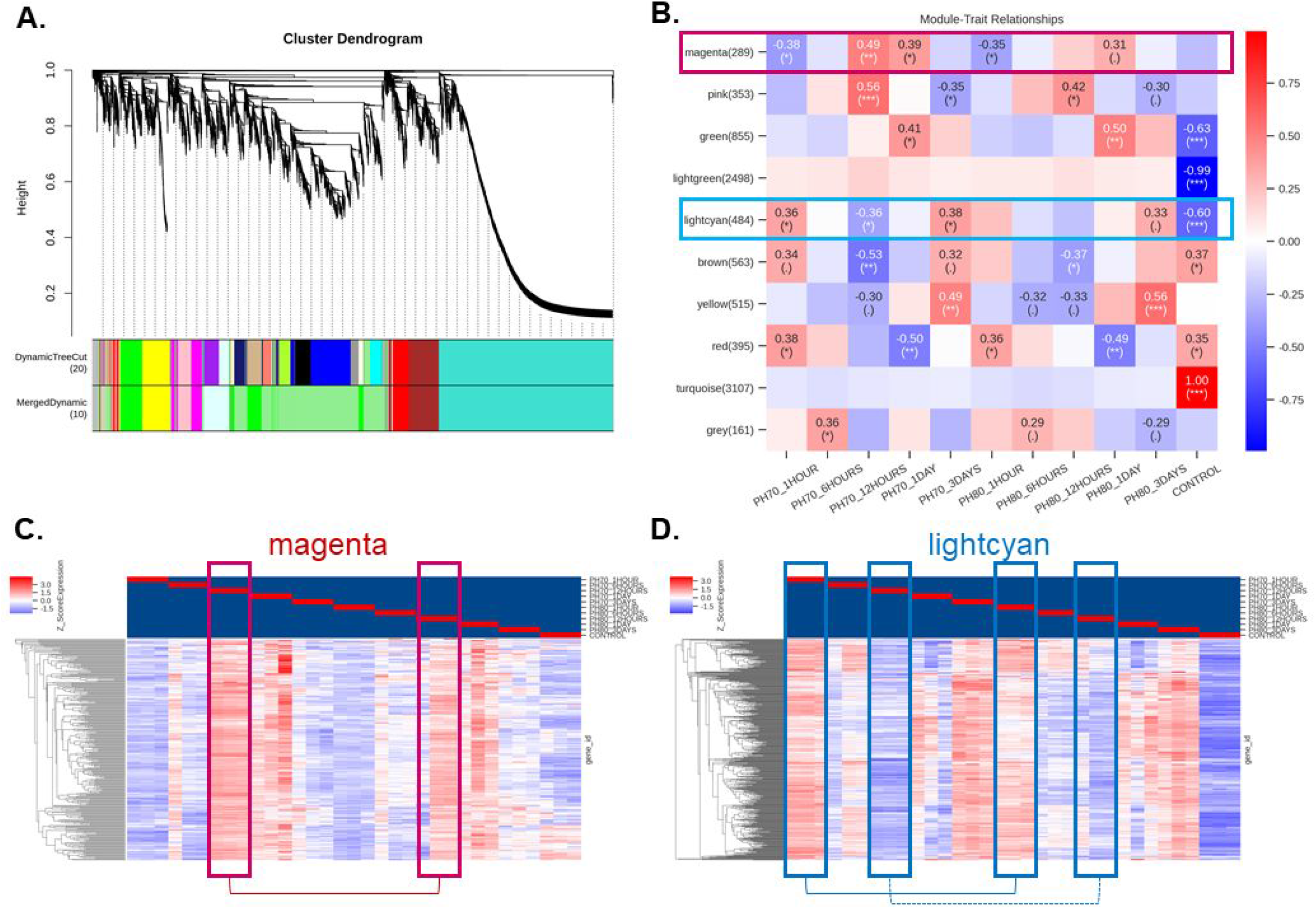
Identification of key modules and module-trait association of hepatectomy by construction of WGCNA. Figure 2A. Module screening based on expression of differentially expressed genes following major hepatectomy and extended hepatectomy. A number of 10 co-expression modules in different colors were constructed. The modules were ranged from large to small by the number of genes included. Figure 2B. Module-trait association of module eigengenes and different time points after hepatectomy. Each row in the table corresponds to a consensus module, and each column to a trait (time point after hepatectomy). Number in the table reports the correlations of the corresponding module eigengenes and traits, with the p-values printed below the correlations in parentheses. Megenta and lightcyan modules are highlighted with color. Figure 2C and 2D. Interaction of 10 co-expression modules using a heatmap plot. Magenta and lightcyan modules in color frame with different association were connected.

The magenta module (gene count: 289) was found positively correlated with major hepatectomy at 12h, and the lightcyan module (gene count: 484) was positively correlated with major hepatectomy at 1h and 12h, while not correlated with extended hepatectomy (Supplemental file 1). The expression profiles of green (gene count: 855), lightgreen (gene count: 2498), pink (gene count: 353), brown (gene count: 563), yellow (gene count: 515), red (gene count: 395) and turquoise (gene count: 3107) modules were similar between major hepatectomy and extended hepatectomy (Figure 1B).

### Functional enrichment analysis of key modules and selection of the differentially expressed gene of key modules in extended hepatectomy

In order to analyze the biological traits of the two key modules, functional enrichment analysis was performed (Supplemental file 2), which were correlated with major hepatectomy not extended hepatectomy. Top 10 GO enrichment and KEGG pathway analyses in the key modules are listed in Table 2 and 3.

**Table 2.** GO enrichment analysis of genes in co-expression modules associated with hepatectomy.

| Module | ID | Term | Category | ListHits | P value | Enrichment_score |
| --- | --- | --- | --- | --- | --- | --- |
| **magenta** | GO:0072378 | blood coagulation, fibrin clot formation | biological_process | 3 | 2.20E-07 | 41.08705 |
|  | GO:0042273 | ribosomal large subunit biogenesis | biological_process | 6 | 6.22E-07 | 11.73916 |
|  | GO:1904874 | positive regulation of telomerase RNA localization to Cajal body | biological_process | 4 | 1.71E-06 | 18.26091 |
|  | GO:0000398 | mRNA splicing, via spliceosome | biological_process | 9 | 4.49E-06 | 5.654181 |
|  | GO:0090277 | positive regulation of peptide hormone secretion | biological_process | 3 | 5.29E-06 | 22.82614 |
|  | GO:0016070 | RNA metabolic process | biological_process | 3 | 8.72E-06 | 20.54353 |
|  | GO:0034622 | cellular protein-containing complex assembly | biological_process | 4 | 1.08E-05 | 13.04351 |
|  | GO:0034116 | positive regulation of heterotypic cell-cell adhesion | biological_process | 3 | 1.35E-05 | 18.67593 |
|  | GO:0006413 | translational initiation | biological_process | 6 | 1.86E-05 | 7.208254 |
|  | GO:0051258 | protein polymerization | biological_process | 3 | 2.01E-05 | 17.1196 |
| **lightcyan** | GO:0050667 | homocysteine metabolic process | biological_process | 4 | 8.41E-07 | 18.80198 |
|  | GO:0033762 | response to glucagon | biological_process | 5 | 1.11E-06 | 13.22014 |
|  | GO:0006563 | L-serine metabolic process | biological_process | 3 | 4.45E-06 | 21.15222 |
|  | GO:0015747 | urate transport | biological_process | 4 | 4.98E-06 | 14.10148 |
|  | GO:1902287 | semaphorin-plexin signaling pathway involved in axon guidance | biological_process | 4 | 4.98E-06 | 14.10148 |
|  | GO:0006820 | anion transport | biological_process | 5 | 9.13E-06 | 9.614646 |
|  | GO:0006629 | lipid metabolic process | biological_process | 10 | 1.82E-05 | 4.361283 |
|  | GO:0055085 | transmembrane transport | biological_process | 17 | 2.85E-05 | 2.888255 |
|  | GO:0005980 | glycogen catabolic process | biological_process | 3 | 3.53E-05 | 14.10148 |
|  | GO:0043401 | steroid hormone mediated signaling pathway | biological_process | 7 | 5.48E-05 | 5.195283 |

**Table 3.**
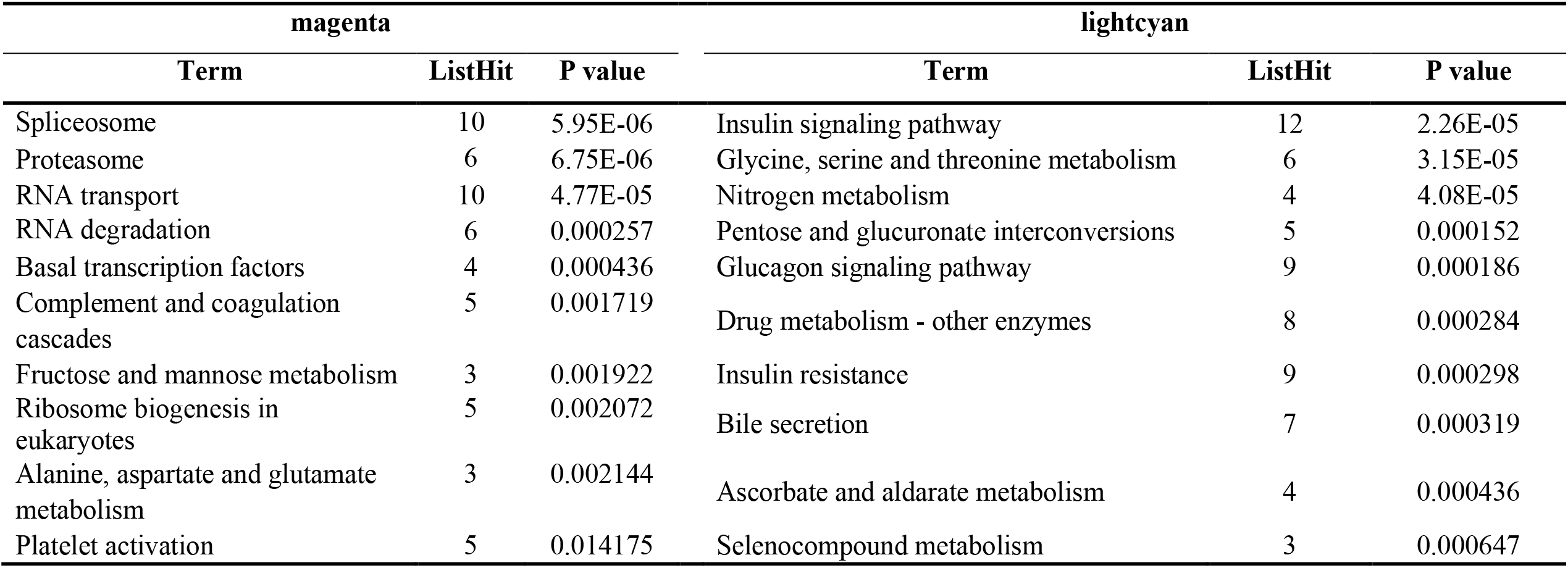
Top 10 KEGG pathways enrichment of modules associated with external traits in hepatectomy.

| magenta |  |  | lightcyan |  |  |
| --- | --- | --- | --- | --- | --- |
| Term | ListHit | P value | Term | ListHit | P value |
| Spliceosome | 10 | 5.95E-06 | Insulin signaling pathway | 12 | 2.26E-05 |
| Proteasome | 6 | 6.75E-06 | Glycine, serine and threonine metabolism | 6 | 3.15E-05 |
| RNA transport | 10 | 4.77E-05 | Nitrogen metabolism | 4 | 4.08E-05 |
| RNA degradation | 6 | 0.000257 | Pentose and glucuronate interconversions | 5 | 0.000152 |
| Basal transcription factors | 4 | 0.000436 | Glucagon signaling pathway | 9 | 0.000186 |
| Complement and coagulation cascades | 5 | 0.001719 | Drug metabolism - other enzymes | 8 | 0.000284 |
| Fructose and mannose metabolism | 3 | 0.001922 | Insulin resistance | 9 | 0.000298 |
| Ribosome biogenesis in eukaryotes | 5 | 0.002072 | Bile secretion | 7 | 0.000319 |
| Alanine, aspartate and glutamate metabolism | 3 | 0.002144 | Ascorbate and aldarate metabolism | 4 | 0.000436 |
| Platelet activation | 5 | 0.014175 | Selenocompound metabolism | 3 | 0.000647 |

The intramodular connectivity of the constructed modules was calculated by summing the connection strengths with other module genes and dividing this number by the maximum intramodular connectivity. The top-50 intramodular hub genes in the key modules were shown in Supplemental Table S3.

To verify the analysis of transcriptome sequencing and WGCNA, PPARα from lightcyan module was selected, and its expression was compared in between PH70 and PH80. Results showed that PPARα expression after extended hepatectomy was significantly lower at 1h after extended hepatectomy in rats by transcriptome sequencing (*t*=8.854, *P*=0.001, Figure 3A), RT-qPCR (*t*=3.006, *P*=0.040, Figure 3B) and WB (*t*=3.840, *P*=0.045, Figure 3C and 3B). Human samples obtained at 1 h after hepatectomy showed PPARα expression after marginal hepatectomy was significantly lower than minor hepatectomy by RT-qPCR (*t*=5.089, *P*=0.001, Figure 3D) and WB (*t*=4.217, *P*=0.001, Figure 3E and 3D).

**Figure 3.**
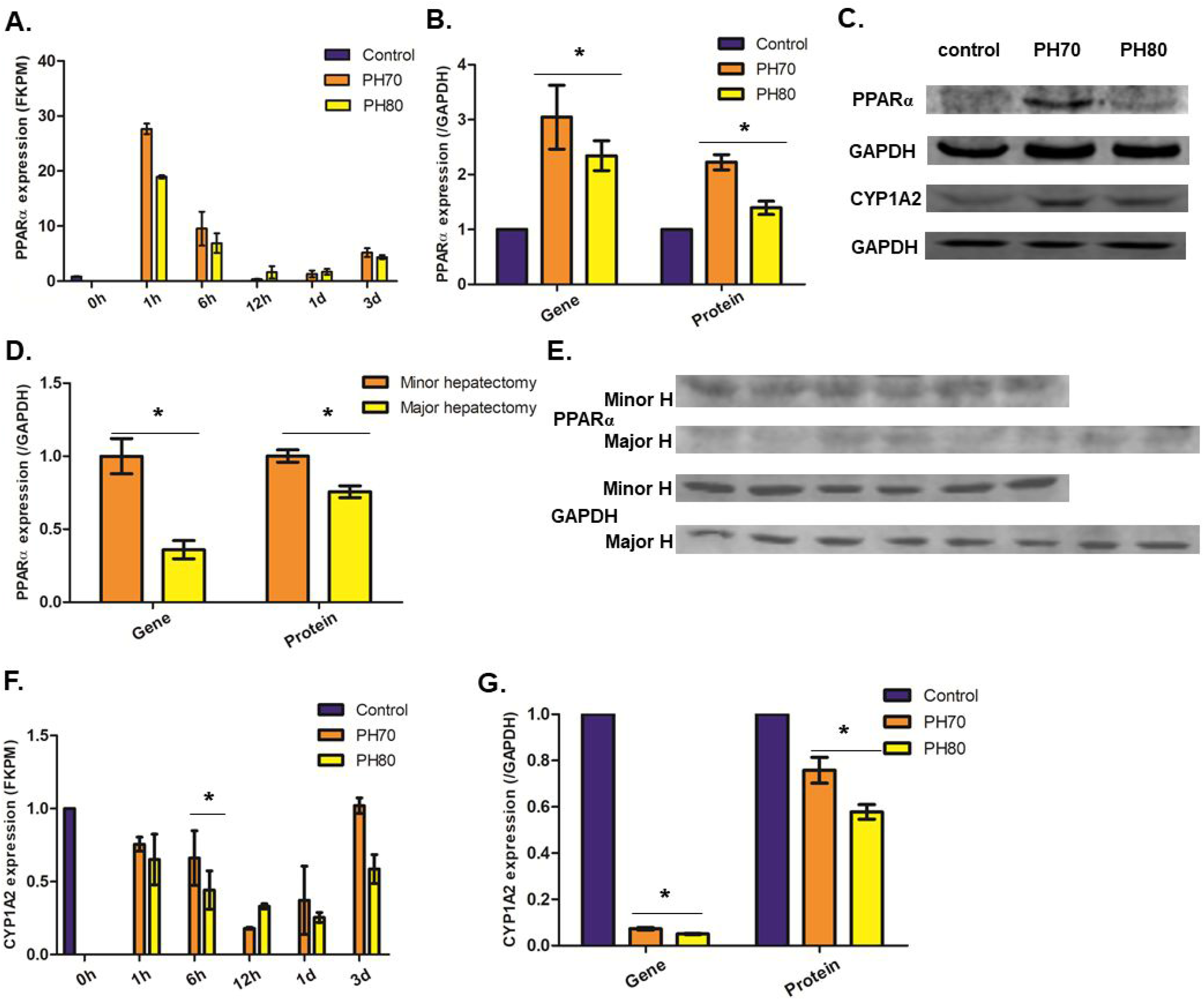
Comparative expression of PPARα following extended hepatectomy. PPARα expressions were compared between major hepatectomy and extended hepatectomy in rats by transcriptome sequencing (Figure 3A), RT-qPCR (Figure 3B) and WB (Figure 3C and 3B). PPARα expressions were compared between marginal hepatectomy and minor hepatectomy in humans by RT-qPCR (Figure 3D) and WB (Figure 3E and 3D). Expressions of PPARα-target gene, CYP2A2, were compared between major hepatectomy and extended hepatectomy in rats by transcriptome sequencing (Figure 3F), RT-qPCR (Figure 3G) and WB (Figure 3C and 3G). Asterisk (*) indicates *P* < 0.05. CYP1A2: cytochrome P450 1A2; PPARα: peroxisome proliferator-activated receptor-α

In addition, PPARα down-stream regulated CYP1A2 was shown significantly lower at 6h after extended hepatectomy in rats by transcriptome sequencing (*t*=12.149, *P*=0.001, Figure 3F), RT-qPCR (*t*=4.089, *P*=0.002, Figure 3G) and WB (*t*=2.788, *P*=0.019, Figure 3C and 3G).

### Functional verification of PPAR*α* in extended hepatectomy

In order to confirm the effect of PPARα, pharmacological activation or inhibition by agonist and antagonist of PPARα was employed to verify the function of PPARα after extended hepatectomy. CYP1A2 was significantly down-regulated after WY14643 administration and upregulated after GW6471 by RT-qPCR (*F*=20.700, *P*=0.001, Figure 4A) and WB (*F*=12.280, *P*=0.001, Figure 4B and 4C).

**Figure 4.**
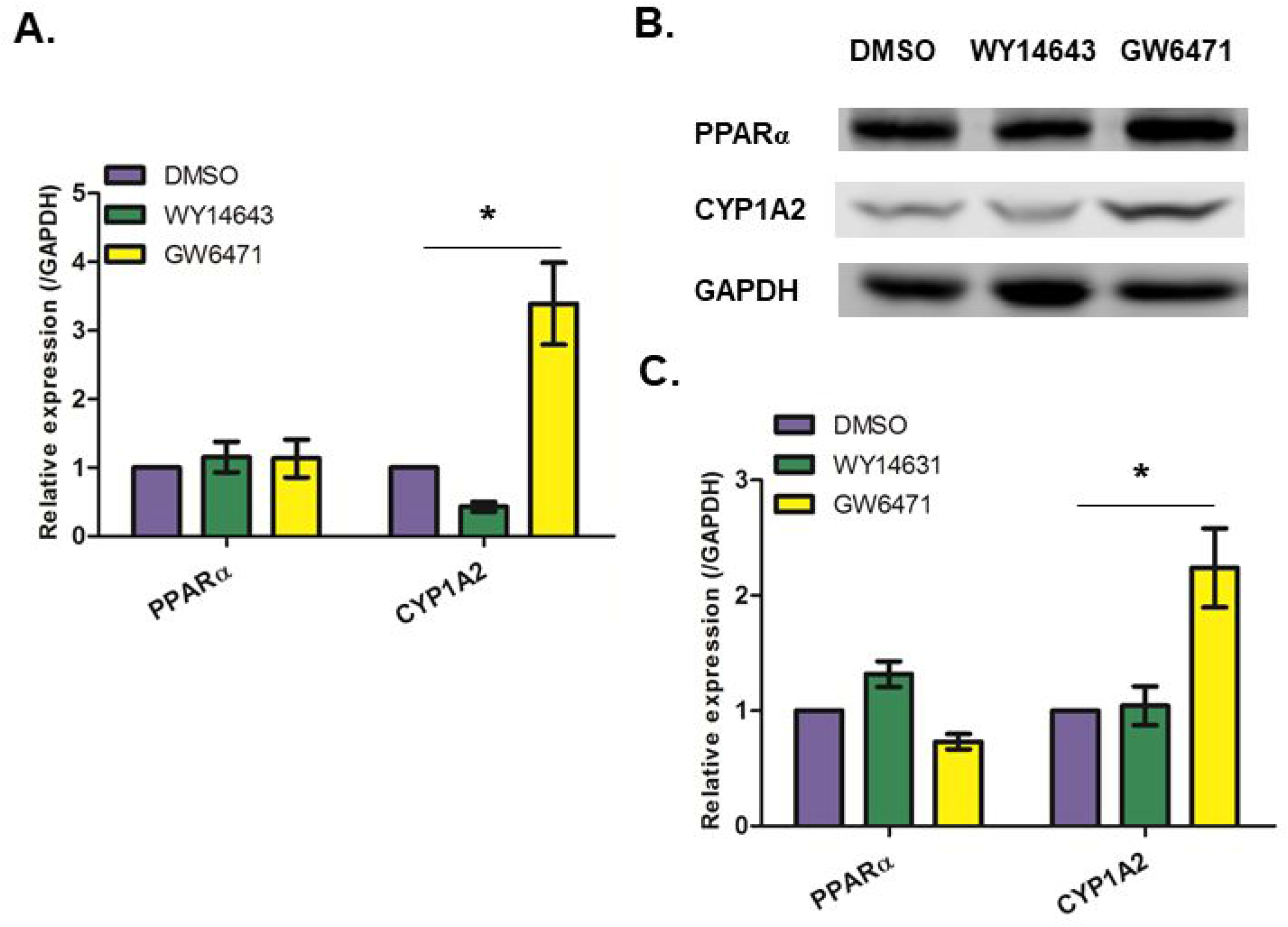
The expressions of PPARα at 1h and CYP1A2 at 6h following extended hepatectomy in rats with administrations of WY14643 and GW6471 by RT-qPCR (Figure 4A) and WB (Figure 4B and 4C) Asterisk (*) indicates *P* < 0.05. CYP1A2: cytochrome P450 1A2; PPARα: peroxisome proliferator-activated receptor-α

Compared with administration of DMSO and PPARα antagonist, PPARα agonist significantly decreased the postoperative levels of ALT (*F*=7.493, *P*=0.027, Figure 5A) and AST on day 1 (*F*=7.851, *P*=0.017, Figure 5B), Suzuki criteria (*F*=10.182 on day 1 and 7.000 on day 3, *P*=0.012 and 0.027 respectively, Figure 5C), TNFα and IL-6 on day 1 (*F*=22.485 and 18.510, *P*=0.001 and 0.001, respectively, Figure 5D and 5E) by ELISA, indicating the protective role of PPARα following extended hepatectomy. Histological changes after administrations of DMSO, WY14643 and GW6471 were visualized with HE staining in the supplemental file (Supplemental Figure S1). The proliferation parameter, LBW, after extended hepatectomy was not significantly changed on day 1 and 3 (*F*=0.108 and 2.381, *P*=0.900 and 0.188, Figure 5F).

**Figure 5.**
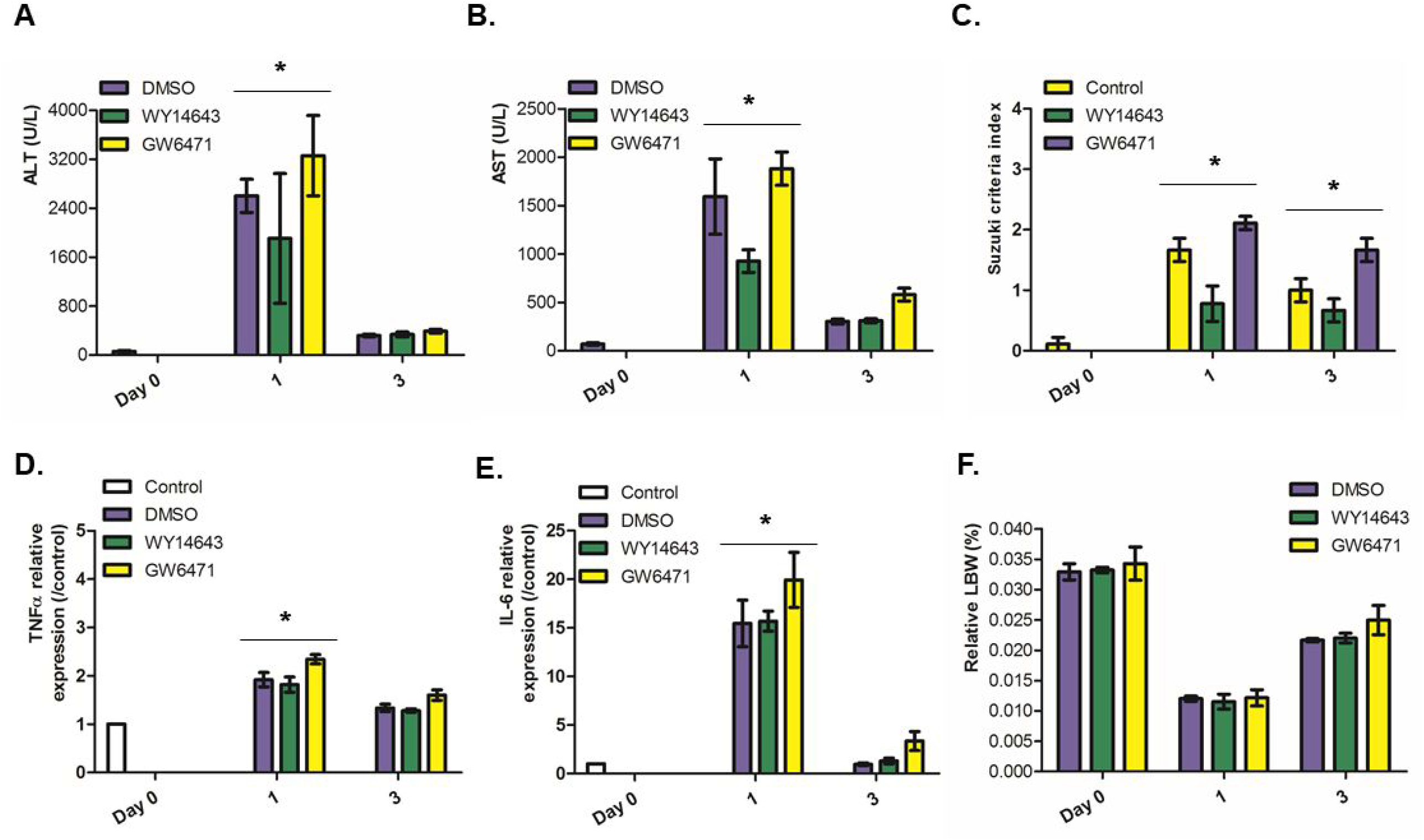
Detection of liver function, injury, inflammation and regeneration following extended hepatectomy in rats and administrations of WY14643 and GW6471. Liver function (ALT and AST, Figure 5A and 5B), injury (Suzuki index, Figure 5C), inflammation markers (TNFα and IL-6, Figure 5D and 5E) and regeneration parameter (LBW, Figure 5F) were compared after administrations of WY14643 and GW6471. Asterisk (*) indicates *P* < 0.05. LBW: remnant liver weight relative to body weight; ALT: alanine aminotransferase; AST: aspartate aminotransferase; TNFα: tumor necrosis factor α; IL-6, interleukin 6

## Discussion

Our previous study showed the 80% extended hepatectomy could stimulate a compromised liver regeneration in comparison with 70% major hepatectomy in rats (Yan et al. 2019), the later which was used as the novel pre-clinic model of liver regeneration. The main focus of the current study was to further compare the transcriptional profiles after between extended hepatectomy and major hepatectomy, and investigate the regulation mechanisms following extended hepatectomy. Through transcriptome sequencing, the differentially expressed genes were identified at the specific time-points following major hepatectomy and extended hepatectomy. Venn diagram illustrated that the number of the differentially expressed genes ranged from 10518 to 11339 at these time points. It was obvious that the differentially expressed genes between major hepatectomy and extended hepatectomy followed the similar expression patterns illustrated in Figure 1. The further bioinformatic and molecular analysis are indicated to identify the “subtle” differences between extended hepatectomy and major hepatectomy. Thereafter, we used WGCNA analysis to outline the molecular profiles and identify the differentially expressed genes from the key modules of magenta and lightcyan. The identified essential genes are indicated as potential biomarkers and targets for intervention. Furthermore, GO and KEGG pathway analysis were conducted to predict the underlying regulation mechanism accounting for the molecular profile after extended hepatectomy.

The magenta module clearly shows enrichment of RNA binding and translation initiation in GO (including molecular function, cellular component, and biological process) and KEGG pathway; in addition, most of genes in lightcyan module show highly enrichment of transporter activity and lipid metabolic process in GO and KEGG pathway analysis. The differentially expressed genes were identified in both magenta and lightcyan modules.

PPARα from lightcyan module was selected to test the bioinformation analysis and to further explore its biological function. PPARα expression was found highly upregulated at the early stage after hepatectomy, while its expression after extended/marginal hepatectomy was found significantly lower in both rats and humans; furthermore, the protective role of PPARα was verified following extended hepatectomy by administration of WY14643 and GW6471, which was in agreement with the previous findings in liver injury (Di Paola and Cuzzocrea 2007; Kersten and Stienstra 2017). Thus, based on transcriptome sequencing and WGCNA analysis after extended hepatectomy and major hepatectomy in rats, the hub genes from key modules were prone for diagnostic biomarker and therapeutic targets for extended/marginal hepatectomy.

To be addressed, the previous study demonstrated that expression of PPARα decreased at 10-26h after partial hepatectomy of rats and mice (Skrtic et al. 2005). Consistent with this finding, our study supported the decreased expression of PPARα at 6-72h following major hepatectomy and extended hepatectomy in rats. More importantly, we observed a rapid increase of PPARα expression at 1-6h after partial hepatectomy in both rats and humans. The up-regulation of PPARα might be related with fatty acid metabolism in response to oxidative response and liver inflammation (Kersten and Stienstra 2017). Besides, regarding to the function of PPARα after major hepatectomy, some studies reported that PPARα may promote liver regeneration (Xie et al. 2019; Brocker et al. 2017). Our study indicated hepatic PPARα influences liver inflammation while not affecting liver regeneration through *in vivo* administration of PPARα agonist and antagonist in the early stage after extended hepatectomy. The mechanisms accounting for the effect of PPARα on liver regeneration after extended hepatectomy are still unclear.

In order to investigate the response and its regulation after partial hepatectomy, a series of previous studies have reported bioinformatic analyses of gene/protein expressions in rodents with a variety of platforms varying from microarray to proteomics (Wu et al. 2010; Zhou et al. 2016; Rychtrmoc et al. 2012). The results varied considerably both in the number and the nature of genes, making the validation difficult. In addition, the clinical application is not mentioned. Our results were based on functional genomic analysis from the experimental models of extended hepatectomy and major hepatectomy. The major weakness of the study is that the biomarker and therapy targets need the further verification in the clinical trials. The regulation mechanisms following extended hepatectomy need more evidences in the further experiments.

In conclusion, the comparative studies of the genomic profile in the current study will help to elucidate the regulation mechanism of post-operative hepatic failure, and thus provides new insights to identify the molecular targets at the transcription level following extended hepatectomy. The selected hub gene, PPARα and PPARα-regulated CYP1A2, might be a suitable biomarker and therapy target for the clinic usage. The conduct of the current study will provide new insights contributing to the clinical beneficiation.

## Acknowledgment

The authors thank Dr. Feng Wang (Department of Pathology, the First Affiliated Hospital of Zhengzhou University, Zhengzhou, China) for the pathological evaluation.

## Funding

The authors sincerely thank the funding from The Medical Science and Technology Program of Henan Province, China (No. SBGJ2018023 and SB201902017) and The National Natural Science Foundation of China (No. 81971881).

## Statements and Declarations

All the authors declare no conflicts of interest and declare that the research was conducted in absence of any commercial or non-financial relationships that could be construed as a potential conflict of interest.

## Availability of data and materials

The datasets generated during and/or analyzed during the current study are available from the corresponding author on reasonable request. The project and the raw data of the transcriptome sequencing have been registered and uploaded in the National Center for Biotechnical Information (PRJNA687124).

## Figure legend

**Supplemental Figure S1.**
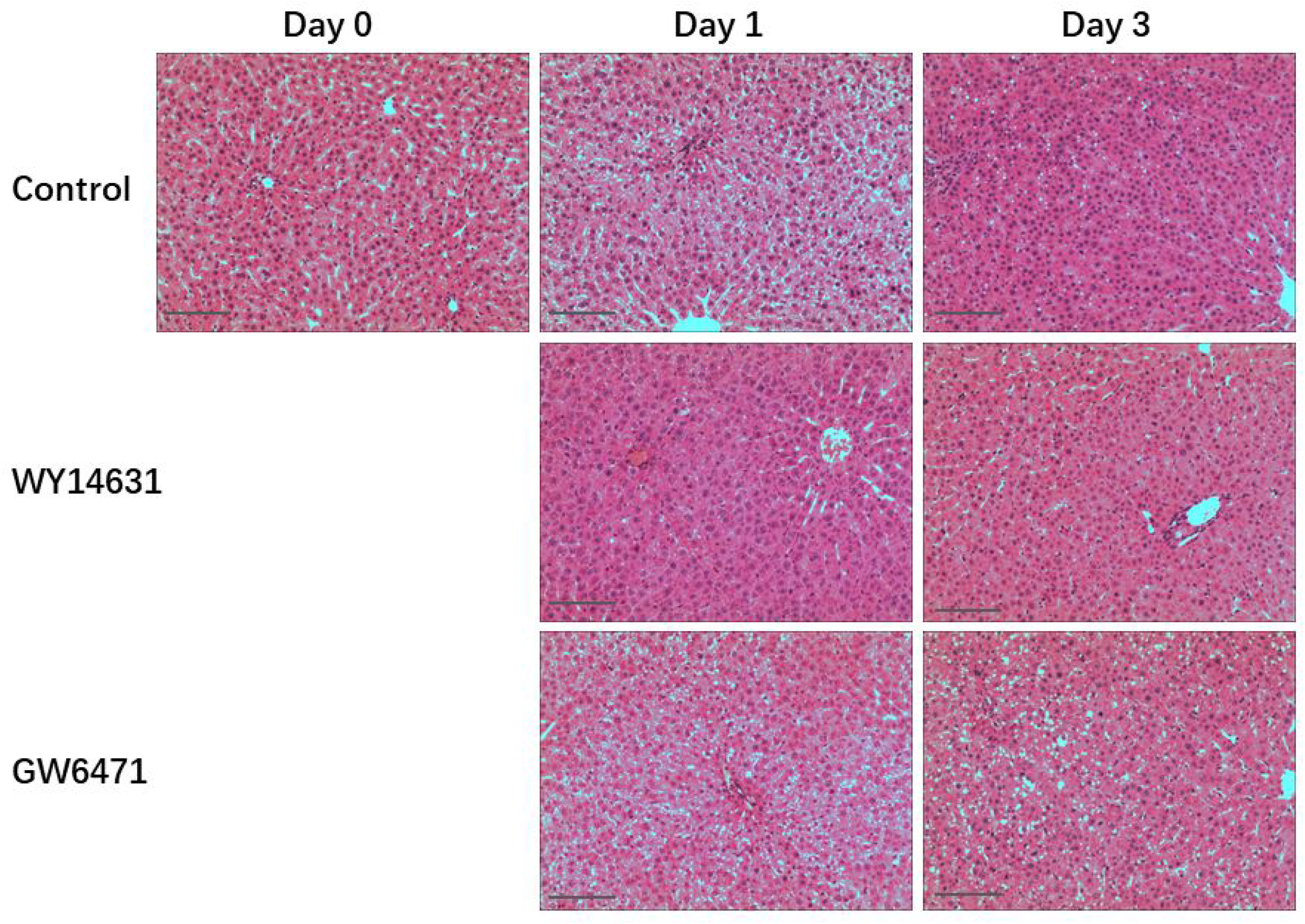
Histological changes after administrations of DMSO, WY14643 and GW6471 were visualized with hematoxylin and eosin staining (hematoxylin counterstaining, original magnification ×200, and scale bars 50μm).

